# A light-dependent molecular link between competition cues and defense responses in plants

**DOI:** 10.1101/637504

**Authors:** G. L. Fernández-Milmanda, C. D. Crocco, M. Reichelt, C. A. Mazza, T. G. Köllner, T. Zhang, M. D. Cargnel, M. Z. Lichy, A. J. Koo, A. T. Austin, J. Gershenzon, C. L. Ballaré

## Abstract

One of the principal internal signals controlling plant growth and defense is jasmonate (JA), a potent growth inhibitor that is simultaneously a central regulator of plant immunity to herbivores and pathogens. When shade-intolerant plants perceive the proximity of competitors using the photoreceptor phytochrome B (phyB), they accelerate growth and down-regulate JA responses. However, the mechanisms by which photoreceptors relay light cues to the JA signaling pathway are not understood. Here we identify a sulfotransferase (ST2a) that is strongly up-regulated by plant proximity perceived by phyB via the phyB-Phytochrome Interacting Factor (PIF) signaling module. By catalyzing the formation of a sulfated JA derivative, ST2a acts to degrade bioactive forms of JA and represents a direct molecular link between photoreceptors and hormone signaling in plants. The enzyme provides a molecular mechanism for prioritizing shade avoidance over defense under close plant competition.

## RESULTS and DISCUSSION

Growth responses to competition with other plants (*1*) and defense responses to the attack of consumer organisms (*2*) are two paradigmatic examples of adaptive phenotypic plasticity in plants. However, the mechanistic and functional links between these responses are not well understood. JAs are potent growth inhibitors (*3*) and regulators of cell division (*4*, *5*), and their role in balancing growth and defense is evolutionarily conserved in land plants, from bryophytes (*6*) to angiosperms (*7*). When shade-intolerant plants perceive a high risk of competition for light with neighboring individuals, they activate the shade-avoidance syndrome (SAS), which allows them to position their leaves in well-illuminated areas of the canopy. Under these competitive conditions, plants also often attenuate the expression of JA-mediated defense responses against pathogens and herbivore (*8*). This attenuation of defense presumably allows the plant to efficiently focus its resources and developmental decisions on escaping shade, sacrificing plant parts that are unlikely to contribute to resource capture.

Plants perceive the proximity of competitors using photoreceptors. Low ratios of red (R) to far-red (FR) radiation (R:FR ratio), which indicate a high risk of competition, result in partial inactivation of the photoreceptor phyB, which in turn promotes growth-related hormonal pathways (*9*), and attenuates signaling mechanisms involved in the activation of defense responses, such as the JA and salicylic acid signaling pathways (*8*). The attenuation of defense responses triggered by supplemental FR radiation or mutations in the *PHYB* gene is not merely a consequence of redirecting resources to growth (i.e., a simple reflection of an energetic tradeoff between growth and defense)(*10*, *11*). Attenuation of JA responses under low R:FR ratios has been associated with FR-induced changes in the balance between DELLA and JA-ZIM-domain (JAZ) repressor proteins (*12*, *13*), and decreased stability of key transcription factors, such as MYC2 (*13*). However, the specific molecular links between the phyB and JA signaling pathways have not been demonstrated.

Most JA-upregulated genes and metabolites are down-regulated by low R:FR ratios or phyB inactivation (*8*, *11*). In order to identify signaling elements that could be involved in the suppression of defense in plants exposed to competition cues, we searched for genes with an inverse pattern of regulation (i.e., JA-related genes showing increased expression under conditions that inactivate phyB). In a microarray experiment, we found a small cluster of approximately 100 genes that were positively regulated by JA whose expression was promoted by phyB inactivation; this cluster included various *JAZ* genes, and a gene of the sulfotransferase family annotated as *ST2a/SOT15* (Fig. S1A). Moerover, in an analysis of expression patterns of JA-related genes using publicly-available microarray databases, we discovered that this *ST2a* gene was consistently and strongly upregulated by low R:FR ratios (Fig. S1B). Experimentally, we confirmed a strong upregulation of *ST2a* mRNA by FR radiation treatments that mimicked the effects of plant proximity and mild suppression by UV-B radiation (Fig. S1C), suggesting that this gene could indeed be involved in relaying information on the canopy light conditions to the JA signaling pathway. Chromatin immunoprecipitation sequencing (ChIP-seq) results indicate that *ST2a* is among the direct targets of Phytochrome Interacting Factors (PIFs) (*14*). PIFs are growth-promoting transcription factors (*15*), and PIF4, PIF5, and PIF7 have been shown to link phyB inactivation with growth responses to shade signals (*16*, *17*). We tested a *pif4 pif5 pif7* triple knock out mutant (*18*) and found that ST2a mRNA was upregulated by tissue damage just like in Col-0, but the transcriptional response of *ST2a* to supplemental FR radiation was completely lost (Fig. S1D). These results demonstrate that low R:FR ratios upregulate the transcription of *ST2a* via the phyB-PIF transcription module.

*ST2a* belongs to a family of 21 sulfotransferase-encoding genes in Arabidopsis (*19*, *20*), and shows sequence similarity to proteins in many dicotyledonous species (Fig. S2). In vitro, the ST2a protein catalyzes the sulfation of 12-hydroxy JA (OH-JA) to form JA sulfate (HSO_4_-JA) (*21*). First described in the late 1800s (*22*), sulfation consists of the transfer of a sulfate residue to a hydroxyl or amino group. In mammalian systems, sulfation represents an important pathway for the biotransformation of hormones, neurotransmitters, and numerous xenobiotics (*23*, *24*), which in most cases results in increased water solubility and decreased biological activity. Based largely on their activities and substrate specificities in vitro, sulfotransferases have also been proposed to be important in the regulation of hormonal signaling in plants (*19*, *20*). For example, in Arabidopsis, a tyrosylprotein-sulfotransferase was reported to regulate the activity of hormone peptides involved in the control of cell proliferation (*25*). However, there is very little evidence from functional genetic studies that sulfotransferases are involved in the adaptive modulation of phytohormonal pathways, and no information about their ecological role in the regulation of plant responses to environmental cues.

We reasoned that a plausible physiological role for the activation of *ST2a* transcription via the phyB-PIF module could be the attenuation of JA-mediated responses in plants exposed to competition signals (low R:FR ratios). To test this hypothesis, we treated Arabidopsis plants with methyl JA (MeJA) and evaluated the formation of JA-related metabolites and accompanying changes in the expression of genes related to JA metabolism. Plants were kept under either white light (Amb light treatment) or white light supplemented with FR radiation (FR treatment). MeJA treatment induced rapid (≤30 min) increases in concentrations of JA and the bioactive conjugate JA-Ile, which were followed by increases in further metabolites, including OH-JA, OH-JA-Ile, and later-on by a marked increase in COOH-JA-Ile and HSO_4_-JA (Fig. 1). Addition of FR to ambient light significantly decreased the abundance of JA and JA-Ile, and their oxidized derivatives. In contrast, the pool of HSO_4_-JA was significantly increased by supplemental FR. The increase in HSO_4_-JA concentration under FR correlated well with a dramatic increase in *ST2a* mRNA level (Fig. S3). Genes encoding other enzymes involved in the metabolism of JAs, such as *IAR3*, *ILL6*, and *CYP94B3* were also up-regulated by FR but to a much lesser extent than *ST2a* (Fig. 1, Fig. S3). In summary, low R:FR ratios decreased the pools of JA and JA-Ile, and this effect coincided with a massive up-regulation of *ST2a* gene expression.

**Fig. 1.**
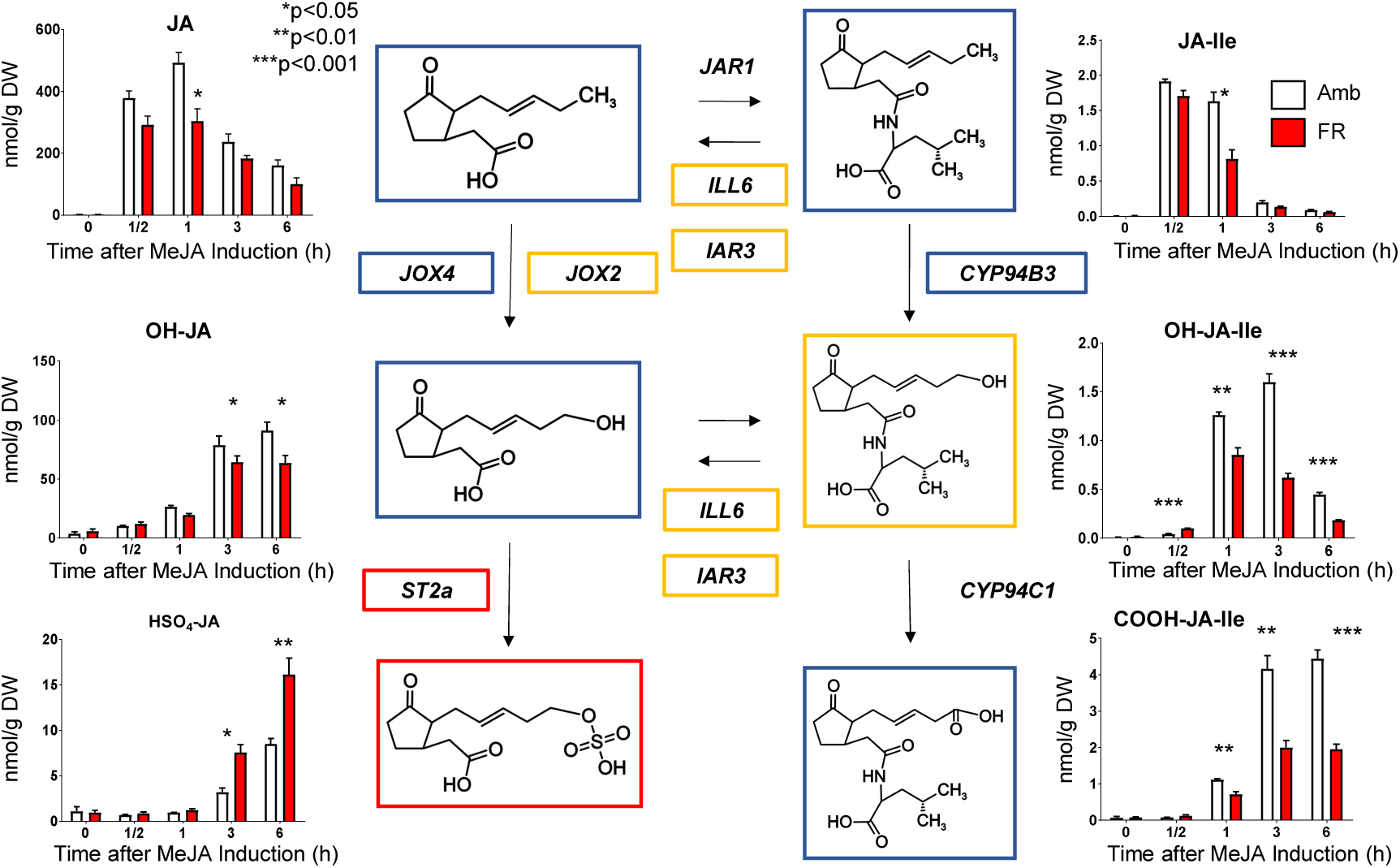
FR supplementation reduces the pool of bioactive JAs and increases *ST2a* transcription and JA sulfation. Col-0 Arabidopsis plants were sprayed with 200 μM MeJA and harvested at the indicated time points for measurements of JA pools and gene expression. The color of the box outline indicates the direction of the FR effect: Blue = downregulation; Yellow = transient upregulation; Red = upregulation; unboxed genes were not significantly regulated by FR. Metabolic map adapted from Wasternack and Feussner (*30*). The bar charts show quantitative data for metabolite concentrations (thin bars indicate 1 SE; n = 3 biological replicates). For gene expression data, see Fig. S3.

In the field, shade treatments, compared to full sunlight, have been reported to attenuate the JA burst induced by insect herbivory, which correlates with attenuated production of defense metabolites (*26*). Are the effects of shade on JA accumulation mediated by phyB inactivation and functionally connected with *ST2a* gene expression? To address this question, we first isolated and characterized two *ST2a* null alleles (*st2a-1* and *st2a-2*), and demonstrated that both knock-out mutants produced only trace levels of HSO_4_-JA (Fig. S4). The mutant carrying the *st2a-1* allele was used for further functional characterization. In Col-0 plants, supplemental FR reduced the JA burst induced by wounding, and this effect correlated with an increase in the pool of HSO_4_-JA (Fig. 2). Importantly, in *st2a-1* plants, the effect of FR attenuating the JA response burst was completely missing (Fig. 2). OH-JA was reduced by FR in Col-0, and it was more abundant in *st2a-1* than in wild type plants (Fig. S5). Within the resolution of our sampling, the levels of JA-Ile were very low and variable, and most of the JA-Ile conjugates were present in the carboxylated form (COOH-JA-Ile) 4 h after wounding. The concentrations of the sum of JA-Ile conjugates was significantly lower in FR plants than in plants of the ambient light treatment (Fig. S5), and significantly higher in *st2a-1* than in Col-0 plants at 4 h. Overaccumulation of COOH-JA-Ile has also been reported in lines lacking JOX/JAOs (*27*), the enzymes responsible for generating OH-JA (the putative substrate of ST2a). Thus, genetic ablation of *JAO/JOX* or *ST2a* results in increased flux through JA-Ile metabolism and catabolite accumulation. Because *ST2a* transcription in response to FR supplementation was minimal in the *pif4 pif5 pif7* triple mutant (Fig. S1D), we used this mutant as a complementary genetic tool to manipulate *ST2a* mRNA levels. Plants of *pif4 pif5 pif7* accumulated HSO_4_-JA in response to wounding, but failed to do so in response to supplemental FR (Fig. S6A). Furthermore, when *ST2a* expression was physiologically manipulated in Col-0 plants using various combinations of FR and wounding treatments, the variation in the HSO_4_-JA content at 4 h after wounding was largely explained by the variation in the levels of *ST2a* mRNA detected by qPCR (Fig. S6B). Finally, when we overexpressed *ST2a* under a strong promoter, we obtained JA metabolite profiles that were qualitatively similar to those generated in response to FR radiation (HSO_4_-JA was greatly increased, with a concomitant reduction in OH-JA, JA, and oxidized JA-Ile conjugates; Fig S7). Neither wounding nor supplemental FR radiation affected the abundance of transcripts of *ST2b*, a gene closely related to *ST2a* (Fig. S2), and a *st2b* null mutant showed normal levels of HSO_4_-JA (Fig. S8). These results provide compelling evidence that ST2a is the sole sulfotransferase responsible for the generation of HSO_4_-JA in Arabidopsis, and that the increased sulfation of OH-JA under low R:FR ratios is functionally connected with the transcriptional upregulation of the *ST2a* gene mediated by the phyB-PIF module. Pharmacological experiments indicate that exogenous applications of OH-JA, the preferred ST2a substrate in vitro, can enhance some JA-Ile triggered responses in Arabidopsis (*27*) and induce the expression of certain genes, including *ST2a* (*21*). Therefore, sulfation of OH-JA by ST2a may be critical for the generation of a genuinely inactive metabolite, channeling JA molecules away from bioactive pools.

**Fig. 2.**
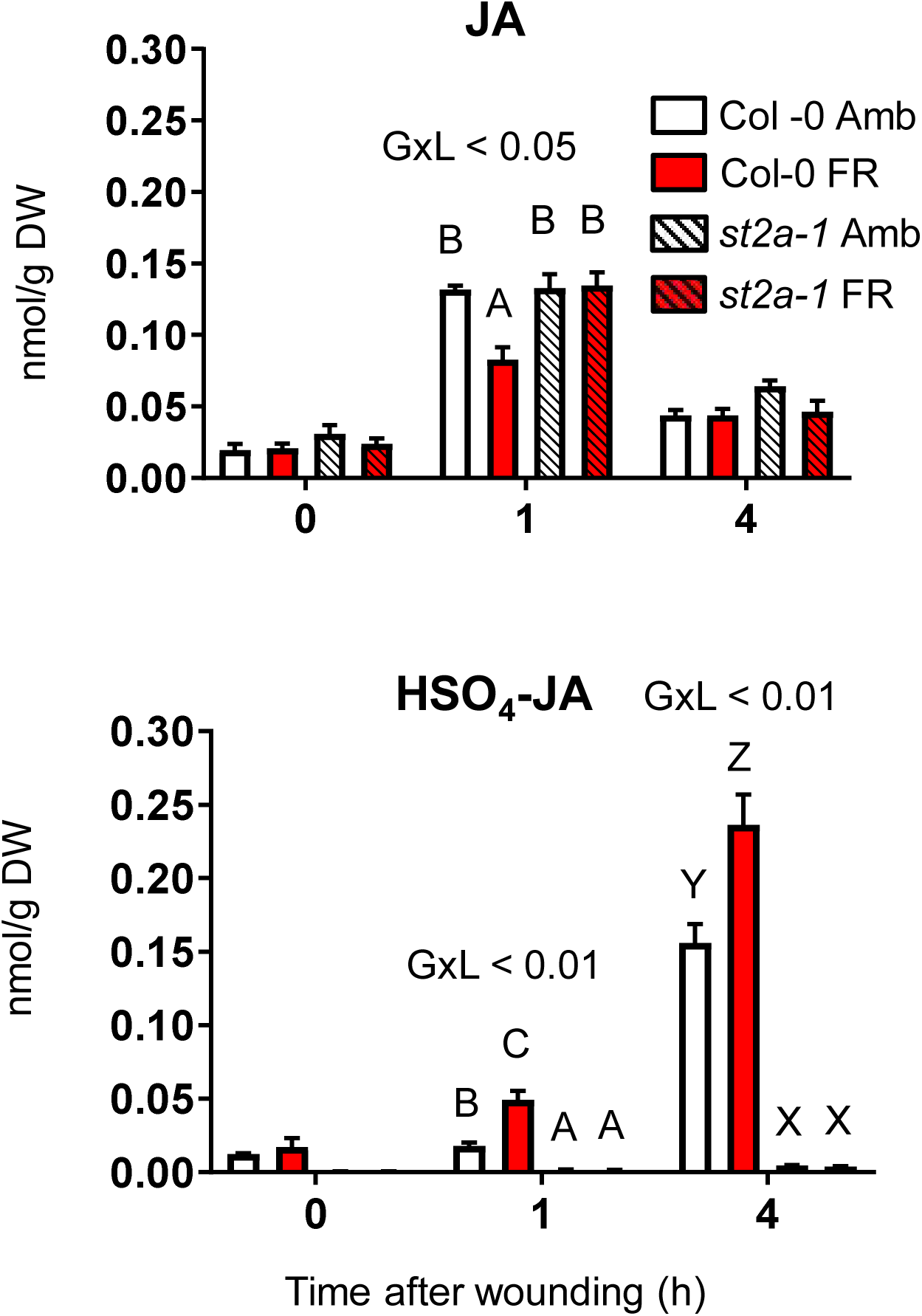
FR attenuates the JA burst triggered by mechanical wounding and increases the concentration of HSO_4_-JA in a *st2a*-dependent manner. Significant genotype x light (GxL) interaction terms are indicted. For each time point, different letters indicate significant differences between means (P < 0.05); thin bars indicate 1 SE (n = 6 biological replicates). DW = dry weight. For additional jasmonate pools, see Fig. S5.

To define the functionality of ST2a, we measured JA-response markers in plants exposed to mechanical wounding under contrasting light conditions. Genes involved in JA biosynthesis (*LOX2*), JA signaling (*MYC2*), and JA response (*VSP2*) were regulated as expected in Col-0, with FR repressing the response to wounding (Fig.3A). In the *st2a-1* null mutant, the basal expression of these genes was higher than in Col-0, and the suppressing effect of FR radiation completely disappeared (Fig. 3A). RNAseq analysis of samples from wounded rosettes revealed a statistically significant overlap between the genes downregulated by FR in Col-0 plants and those upregulated by the *ST2a-1* mutation under FR radiation. The group of overlapping genes was significantly enriched in the GO terms “Response to JA” and “JA biosynthetic process” (Fig. 3B) and Data File S1). Consistent with the pattern of expression of JA biosynthetic genes in Col-0 and *ST2a-1*, we found that FR reduced the accumulation of *cis*-12-oxo-phytodienoic acid (*cis*-OPDA) in Col-0, particularly at high rates of FR supplementation, but this effect of FR was less marked in *ST2a-1* plants (Fig. S9). Glucosinolates (GS) are important defense compounds in Arabidopsis, which are often regulated by JA (*28*) (Fig. S10). In Col-0, the accumulation of these JA-dependent compounds was attenuated when plants were exposed to supplemental FR radiation (Fig. 3C), as expected (*29*). In contrast, in *ST2a-1* plants, FR failed to inhibit GS accumulation (Fig. 3C). Collectively, these data (Fig. 3) indicate that the sulfation reaction catalyzed by ST2a plays a central role suppressing JA-dependent responses in plants undergoing shade avoidance.

**Fig. 3.**
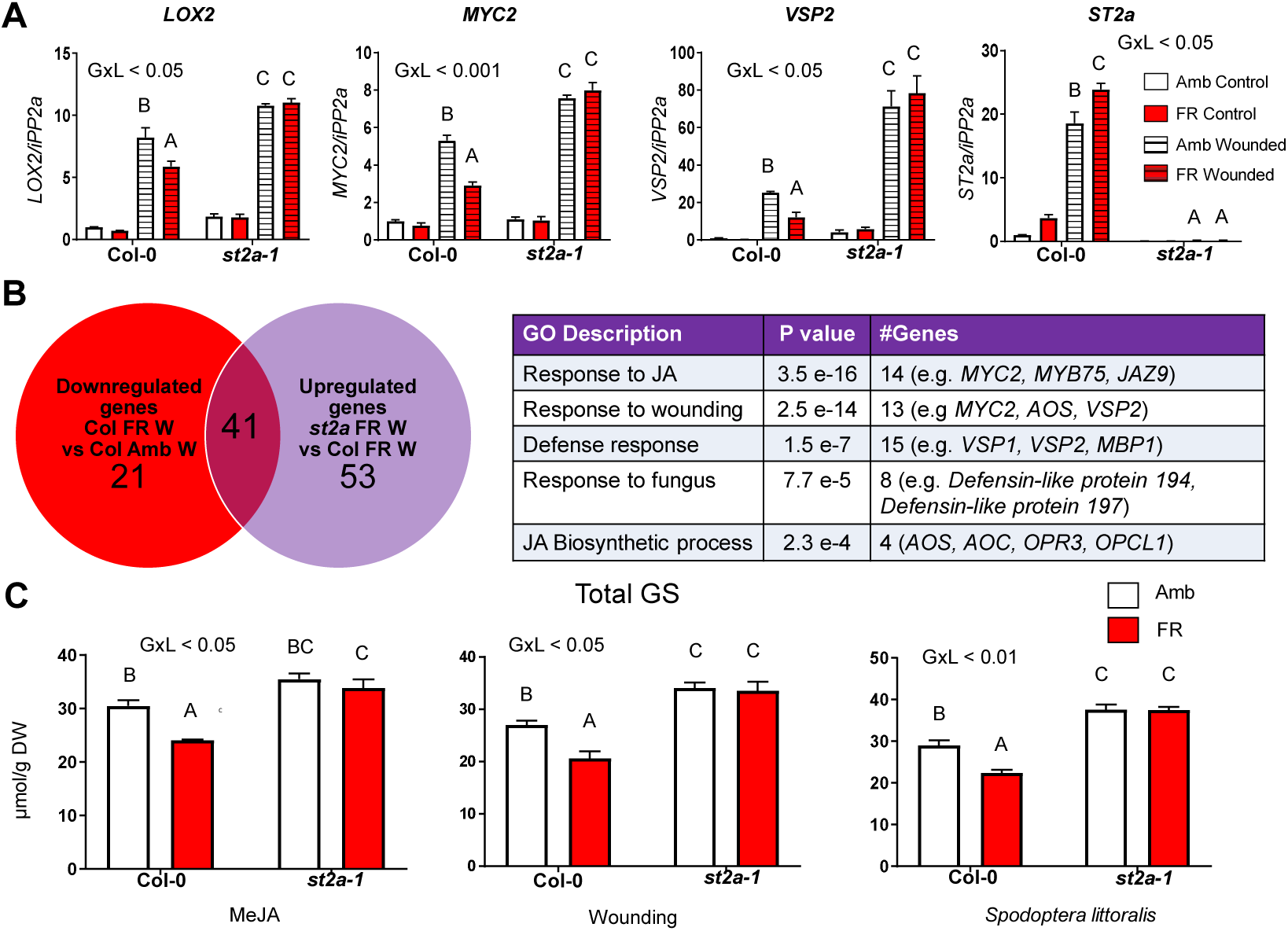
FR downregulates gene and metabolite markers of jasmonate signaling in an *ST2a*-dependent manner. (A) qPCR results for selected markers of JA synthesis, signaling and response. (B) Summary of RNAseq results demonstrating a significant overlap between the genes downregulated by FR in Col-0 plants and those upregulated by the *ST2a-1* mutation in wounded plants. The table shows the GO categories overrepresented in the set of overlapping genes (for details on analysis see Data File S1). (C) Suppression by FR of glucosinolate accumulation in plants treated with MeJA, mechanical wounding or insect herbivory (Spodoptera littoralis) was missing in a *st2a-1* null mutant. For specific data on 4MSOB and I3M in wounded plants, see Fig. S10 B). For induced plants, the significance of the genotype x light (GxL) interaction term is indicated in panels A and C. Different letters indicate significant (P < 0.05) differences between means; thin bars indicate 1 SE (n = 6 biological replicates for glucosinolate data o 3 for transcriptomic data).

To investigate the functional role of changes in JA metabolism caused by *ST2a* activity, we tested the *ST2a-1* null mutant in bioassays with larvae of *Spodoptera littoralis* (a chewing insect) and *Botrytis cinerea* (a necrotrophic pathogen). In Col-0, supplemental FR radiation caused increased growth of *S. littoralis* caterpillars that fed on the plants, and increased the size of necrotic lesions generated by *B. cinerea* (Fig. 4A). These FR effects were missing in plants of *ST2a-1*, which correlated strongly with the lack of effect of FR reducing the concentration of JA (Fig. 2), JA marker gene transcripts, and defense compounds (Fig. 3). Furthermore, the *pif4 pif5 pif7* triple mutant, which did not upregulate the transcription of *ST2a* in response to supplemental FR, was significantly more resistant to *B. cinerea* than Col-0 under low R:FR ratios (Fig. 4A). These data provide compelling empirical support for a functional connection between *ST2a* transcription, increased JA catabolism, and reduced defense under low R:FR ratios.

Rosettes of the *ST2a-1* null mutant appeared similar to those of Col-0 under ambient light, and they showed normal morphological responses to supplemental FR radiation (leaf hyponasty and petiole elongation) (Fig. 4B). However, when plants were treated with MeJA, the shade-avoidance response to supplemental FR radiation was significantly attenuated in *ST2a-1*; in contrast, Col-0 plants were capable of reconfiguring their morphology, and showed a normal response to FR even under MeJA elicitation (Fig. 4B) and Fig. S11). Taken together, these results suggest that the key function of the sulfotransferase ST2a under low R:FR ratios is to facilitate the inactivation of JA, thereby allowing the plant to express its full repertoire of shade-avoidance responses and maximize its competitive ability in crowded stands.

**Fig. 4.**
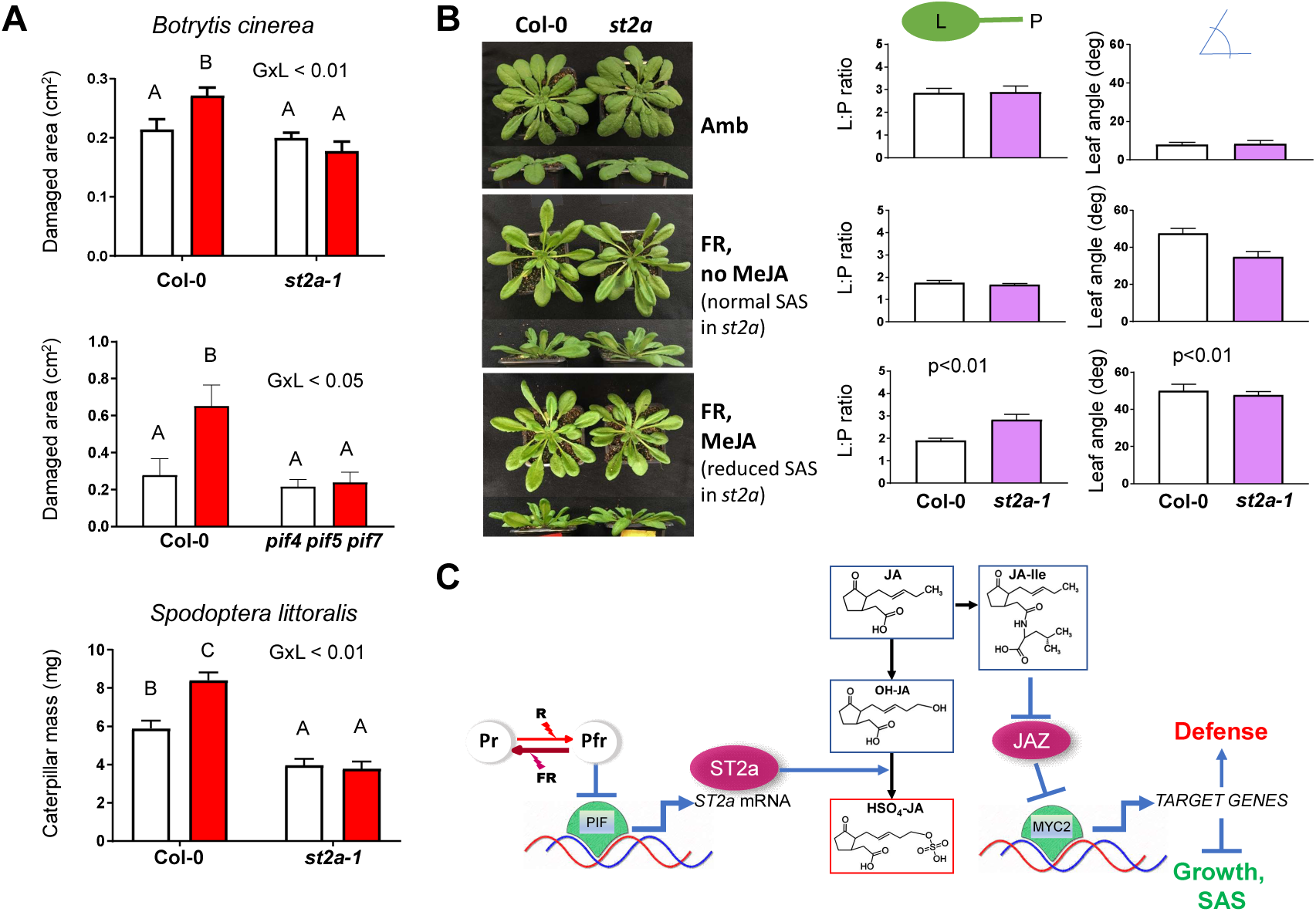
Sulfotransferase *ST2a* is key in regulating the growth/defense balance in response to changes in the R:FR ratio. (A) Under FR supplementation, plants that do not upregulate *ST2a* expression are better defended than their Col-0 counterparts. Bioassays were carried out comparing Col-0 and *st2a-1* rosettes using larvae of *Spodoptera littoralis* (upper panel) and inoculations with *Botrytis cinerea* spore suspensions (middle panel), and also comparing Col-0 and *pif4pif5pif7* triple mutants inoculated with *B. cinerea* spore suspensions (lower panel). The significance of the genotype x light interaction term (GxL) is indicated for each factorial experiment; different letters indicate significant (p< 0.05) differences between means. (B) *ST2a-1* rosettes display normal phenotypes under control conditions but, compared with Col-0 rosettes, they display impaired shade-avoidance responses when exposed to low doses of MeJA (100 μM). **, p<0.01; for full dataset, see Fig. S10. (C) Conceptual model linking the perception of low R:FR ratios via phyB with the modulation of jasmonate metabolism and signaling through regulation of *ST2a* transcription via the phyB-PIF transcription module. Pr, inactive form of phytochrome; Pfr, active form of phytochrome.

Failure to respond to competition signals with a rapid reconfiguration of shoot architecture and leaf traits carries a disproportionate fitness penalty for plants competing for light in fast growing stands. Under these conditions, suppression of the ‘growth brake’ (*3*, *4*) imposed by JA could be a key determinant of success, even if it comes at the cost of attenuating defense responses. Our results demonstrate the molecular mechanism that links neighbor perception via phyB with the attenuation of JA signaling (Fig. 4C), and provide a compelling example of the role of sulfotransferases in the adaptive modulation of hormonal metabolism in plants. This phyB-dependent sulfation mechanism generates a metabolic sink for bioactive JA, and allows the plant to refocus its strategy on rapid growth when the perceived risk of competition for light is strong.

## ACKNOWLEDGMENTS

We thank I. Cerrudo, P. Karssemeijer and B. F. Alliani for help with the initial characterization of *ST2a* lines and P. V. Demkura and B. Rothe for technical assistance.

## Author contributions

G.L.F.-M. contributed to all aspects of this research; C.D.C. contributed to experimental design, data collection, analysis and interpretation; M.R. designed and executed protocols for metabolite and hormone analyses; T.Z. and A.J.K. designed and performed hormone profiling; M.D.C. carried out bioassays and M.Z.L. screened mutant lines and helped with gene expression analyses; C.A.M. and T.G.K. performed analyses of transcriptomic data and helped with data interpretation; A.T.A. and J.G. contributed to the general conception of the project and data interpretation; C.L.B. conceived the project and contributed to data generation and analysis, and wrote manuscript with input from all co-authors.

## Data and materials availability

All data needed to evaluate the conclusions in the paper are present in the main text or the supplementary materials.

