## Supplementary Information for "A light-dependent molecular link between competition cues and defense responses in plants"

[1]IFEVA, Consejo Nacional de Investigaciones Científicas y Técnicas – Universidad de Buenos Aires, Ave. San Martín 4453, C1417DSE, Buenos Aires, Argentina

[2]Max Planck Institute for Chemical Ecology, Hans-Knöll-Str. 8, 07745 Jena, Germany

[3]Department of Biochemistry, University of Missouri, Columbia 65211, USA

[4]IIBIO, Consejo Nacional de Investigaciones Científicas y Técnicas–Universidad Nacional de San Martín, B1650HMP Buenos Aires, Argentina

[5]Current address: College of Agriculture, South China Agricultural University, Guangdong, China

**This PDF file includes:**

**Materials and Methods**
**Figs. S1 to S11**
**Tables S1 to S3**
**References**
**See attached Data Files S1 and S2**

**Materials and Methods**

Plant material and growth conditions

The following *Arabidopsis thaliana* null mutants were used: *st2a-1* (GABI_149G04), *st2a-2* (SALK_075656)*, st2b* (SALK_009093)*, pif4 pif5 pif7*, *aos*, and *phyB-9.* SALK lines were obtained from the Nottingham Arabidopsis Stock Centre (NASC) and checked by PCR. Seeds of the *pif4pif5pif7* triple mutant (*18*) were provided by Prof. Christian Fankhauser. Seeds of the *aos* null mutant (*31*) (in Col-0 background) were provided by Prof. Edward Farmer. Homozygous *st2a-2* plants were obtained by selection after genotyping the heterozygous seed batch included in NASC stock number N575656. *ST2aOE* lines were generated by PCR-amplifying the ORF of *ST2a* with a primer pair (5'-GCT CTA GAA TGG CTA CCT CAA GCA TGA AG-3' and 5'- GCG TCG ACT TAG CTC AAC CTG AAA GTG-3') and cloning into a plant binary vector pBITS (*32*) at the XbaI and SalI site which places the gene under the control of the cauliflower mosaic virus 35S promoter. Seeds were directly placed in 7 x 7 x 8-cm pots containing a standard substrate mix (80 % Fruhstorfer Nullerde pH = 6.0-6.5, 10 % vermiculite and 10 % sand). Soil was fertilized with 1 g Triabon 3-4 M (Mehrnährstoffdünger 16+8+12)/L soil, 1 g Osmocote Exact Mini 3-4 M (16:8:11)/L soil and watered with a suspension of *Steinernema feltiae* (Katz Biotech AG, Germany). Seeds were stratified in the dark for 3-4 days at 4°C and then placed in a growth chamber under short days (10 h light/14 h dark, 18-20 °C, humidity 50-60 %) and 110 µmol $s^{-1}$ $m^{-2}$ of photosynthetically active radiation (PAR) provided by fluorescent bulbs. Rosette-stage plants of similar age (3-4 weeks old) and size were selected for experiments and randomly assigned to treatments.

Light treatments

We tested the effect of FR supplementation on gene expression, oxylipin and glucosinolate chemistry, growth and defense. We compared two light conditions. Our control condition was white light provided from fluorescent bulbs (hereafter called "Ambient = Amb") (Fig. S11). Our FR supplementation treatment (hereafter called FR) was designed to mimic the effect of FR radiation reflected by neighboring plants in a canopy with a leaf area index <1 (*33*). The plants in the FR treatment were exposed to the same amount of photosynthetically active radiation (PAR) as the plants of the Amb control (110 µmol $s^{-1}$ $m^{-2}$) but received FR radiation from one side during the course of the photoperiod. FR (see spectrum on Fig. S11) was provided by custom-made 120-cm long linear arrays of 5-Watt FR740-05-00-00 LEDs (Roschwege GmbH), and filtered through one layer of a blue polyester film (Roscolux, Supergel, Cinegel #83 Medium Blue) to remove residual red light emitted by the LED units. Except indicated otherwise, the plants received 28 µmol $m^{-2}$ $s^{-1}$ of lateral FR. Other fluence rates of FR supplementation were obtained by adjusting the LED output with an electronic dimmer. The FR supplementation treatment started 1 h before the end of the photoperiod of the day that preceded the start of the wounding treatments, MeJA applications or initiation of feeding or infection bioassays. Spectral scans were obtained with a calibrated Ocean Optics USB4000 micro-spectroradiometer and SpectraSuite analysis software (Ocean Optics).

For the UV-B experiment reported in Fig. S1, plants were exposed to either Amb light or Amb light supplemented with 0.4 µmol $m^{-2}$ $s^{-1}$ of UV-B radiation [measured with a cosine corrected UVB detector (SUD/240/W) attached to a IL-1700 research radiometer (International Light, USA)] provided by a Philips TL/01 UV bulb. Daily applications of this UV-B treatment reduced rosette diameter of Col-0 plants by c.

30 %, without significantly affecting the area growth of rosettes of the *uvr8-6* null mutant (Carlos Crocco and Ana Medina, unpublished data).

Methyl jasmonate (MeJA) treatment and wounding experiments

For induction with MeJA, we sprayed seedlings with the indicated concentrations of MeJA or mock solutions (*34*). Plant responses to mechanical wounding were assessed by gently pressing with a forceps every petiolated leaf of the rosettes (one wound per leaf). In one experiment in which we tested the effects of different wounding intensities on gene expression and metabolite accumulation (Fig. S7), we used the teeth of plastic comb to inflict different amounts of damage to the rosette leaves. Plants were harvested at the indicated time points after treatment and immediately frozen in liquid nitrogen. Samples were stored at -80°C.

DNA extraction and genotyping

To test the NASC lines, seeds were germinated and leaves form individual plants were collected and used for DNA extraction. Leaf tissue was homogenized on EDM buffer and centrifuged at full speed for 5 min, to pellet cellular debris. A fraction of 300 µL of the supernatant was mixed with equal volume of isopropanol and centrifuged at max speed for DNA precipitation. The pellet was dried and resuspended in 100 µL of MilliQ water. PCR was performed using Pfu polymerase (PB-L, Argentina) according to the manufacturer's instructions, with 1 µL of DNA solution and primers to final concentration of 1 µM. Primers were designed to amplify on regions adjacent to T-DNA insertion and their sequences are listed on Table S1.

Gene expression analyses

Total RNA was extracted using a Spectrum Plant Total RNA kit (Sigma Aldrich). The remaining DNA was eliminated from samples using a TURBO DNA-free kit (ThermoFisher).

For quantitative real time PCR (qPCR) analysis, cDNAs were obtained with Superscript III Reverse Transcriptase (Invitrogen, USA) following the manufacturer's instructions. qPCR was performed with 7500 real-time (Applied Biosystems) or CFX Connect real-time (Bio-Rad) qPCR systems following standard methods using SsoAdvanced Universal SYBR Green Supermix (BIO-RAD) and primers to final concentration of 500 nM (annealing temperature 60°C). *IPP2a*, *UBC* or *ACT8* genes were used to normalize for different concentrations of cDNA as their expression was not affected by our treatments and did not vary among the different genotypes. The relative expression levels were calculated using the $2^{-\Delta\Delta Ct}$ method (n=3 pools of three individual plants). Primer sequences are listed in Table S1.

For microarray analyses contrasting Col-0 and *phyB* plants, we used a factorial experimental design with two genotypes (Col-0 and *phyb-9*) and two treatments (MeJA or mock). 4-week old plants were sprayed with MeJA (200 µM) or mock solution and harvested 3 h later. Two biological replicates per treatment were performed for Col-0 and 3 replicates for *phyb-9* (each replicate was a pool of 4 individual plants) RNA was prepared, labelled and hybridized to the arrays in accordance with the manufacturer's instructions for the GeneChip Arabidopsis Gene 1.0 ST Array (Affymetrix). Raw microarray data were processed and analyzed using Affymetrix Expression Console® software. Genes with 'absent' calls and a signal of <50 units in all replicate experiments were filtered out. Significantly differentially expressed genes were identified by performing profile analysis using Significance Analysis of Microarrays (*35*) with a p-value <0.05. A test filter was performed to work only with those genes for which the ratio of expression showed at least a 2-fold change between “Col mock” vs “Col MeJA”. Clusters were generated using DNA-Chip Analyzer (dChip) (*36*). Raw reads were deposited in the NCBI Gene Expression Omnibus (GEO) under the accession (*will be inserted later*).

For RNAseq analysis of Col-0 and *st2a* plants, we used a factorial combination of 3 genotypes, 2 light treatments (Amb and FR), and 2 harvest times (0 and 4 h after wounding). There were 3 biological replicates for each combination, each consisting of a pool of 3 individual rosettes. TruSeq RNA-compatible libraries were prepared from DNAse-treated total RNA and PolyA enrichment was performed before sequencing the transcriptomes on an IlluminaHiSeq 3000 sequencer (Max-Planck-Genome-centre, Cologne, Germany) with 20 Mio reads per library, 150 base pair, single end. Trimming of the obtained Illumina reads and mapping to the Arabidopsis gene model version Araport 11 (https://www.araport.org/data/araport11; primary transcripts) were performed with the program CLC Genomics Workbench (Qiagen Bioinformatics) (mapping parameter: length fraction, 0.8; similarity fraction, 0.9; max number of hits, 15). Empirical analysis of digital gene expression (EDGE) implemented in the program CLC Genomics Workbench was used for gene expression analysis. Raw reads were deposited in the NCBI Sequence Read Archive (SRA) under the accession (*will be inserted later*).

Microarray data mining and GO enrichment analysis

Mining of gene expression data in publicly available databases was performed using the set or microarray experiments described in Mazza et al. (*37*) (for Fig S1B) and GENEVESTIGATOR (*38*). Gene ontology (GO) analysis of enriched functional categories (For Fig. 3) was performed using PANTHER (v14.0) (*39*) for GO aspect biological process and *Arabidopsis thaliana* as organism.

Oxylipin analyses

Except indicated otherwise, we used 6 biological replicates (each consisting of 3 individual rosettes) for each genotype and treatment combination. About 20 mg of lyophilized or 200 mg of fresh *A. thaliana* leaf tissue were homogenized and extracted in 1 ml methanol containing 40 ng D4-SA (Santa Cruz Biotechnology, USA), 40 ng D6-JA (HPC Standards GmbH, Germany), 40 ng D6-ABA (Santa Cruz Biotechnology, USA), and 8 ng D6-JA-Ile (HPC Standards GmbH) as internal standards. Samples were agitated on a horizontal shaker at room temperature for 10 min, and then centrifuged at 14000 rpm for 10 min. An aliquot of the extract was transferred to a 96 Deepwell plate and directly used for phytohormone analysis.

Phytohormone analysis was performed by LC-MS/MS as in Vadassery et al. (*40*) on an Agilent 1260 series HPLC system (Agilent Technologies) with the modification that a tandem mass spectrometer QTRAP 6500 (SCIEX, Darmstadt, Germany) was used. Since we observed that both, the D6-labeled JA and D6-labeled JA-Ile standards (HPC Standards GmbH, Cunnersdorf, Germany) contained 40 % of the corresponding D5-labeled compounds, the sum of the peak areas of D5- and D6-compound was used for quantification. Details of the instrument parameters and response factors for quantification can be found in Table S2. Alternatively, for the results reported in Fig. S7, JA metabolite content was quantified using ultra performance liquid chromatography tandem mass spectrometry (UPLC-MS/MS) (Acquity/Xevo TQ-S system, Waters) based on methods described previously (*41*). Characteristic MS transitions were monitored using multiple reaction monitoring in ESI negative mode for JA (*m/z*, 209 → 59), dihydroJA (211 → 59), OH-JA (225 → 59), $HSO_4$-JA (305 → 97), JA-Ile (322 → 130), [$^{13}C_6$]-JA-Ile (328

→ 136), OH-JA-Ile (338 → 130), and COOH-JA-Ile (352 → 130). Data analysis was carried out using MassLynx 4.1 and TargetLynx software (Waters).

Glucosinolate (GS) analysis by HPLC-UV

Unless otherwise indicated, we used 6 biological replicates (each consisting of 3 individual rosettes) for each genotype and treatment combination. Samples were freeze dried to a constant mass and ground to a fine powder. 20 mg of freeze-dried and pulverized material per sample was used for GS analysis. GS were extracted with 1 mL of 80 % methanol solution containing 0.05 mM intact 4-hydroxybenzylglucosinolate as internal standard. Samples were shaken on a horizontal shaker at room temperature for 10 min, and then centrifuged at 14000 rpm for 10 min. Next, extracts were loaded onto DEAE Sephadex A 25 columns (Sigma–Aldrich) column and washed with 80 % methanol solution, water, and 0.02M MES buffer (pH 5.2). Sulfatase solution (arylsulfatase from Sigma-Aldrich) was applied on the column and incubated at room temperature overnight. Distilled water (500 µl) was used to elute the desulfo-GSs into 96-deep well plates for HPLC-UV analysis (*42*).

The eluted desulfo-GSs were separated using high performance liquid chromatography (Agilent 1100 HPLC system, Agilent Technologies) on a reversed phase C-18 column (Nucleodur Sphinx RP; 250 x 4.6 mm, 5µm particle size; Macherey-Nagel, Düren, Germany) with an water-acetonitrile gradient (1.5% acetonitrile for 1 min, 1.5 to 5% acetonitrile from 1 to 6 min, 5 to 7% acetonitrile from 6 to 8 min, 7 to 21% acetonitrile from 8 to 18 min, 21 to 29 % acetonitrile from 18 to 23 min, followed by a washing cycle; flow 1.0 mL min-1). Detection was performed with a photodiode array detector and peaks were integrated at 229 nm. We used the following response factors: aliphatic GS 2.0, indole GS 0.5, for quantification of individual GSs (11). The following GSs were quantified: 4-methylsulfinylbutyl GS (4MSOB), indol-3-methylsulfinylpropyl GS (3MSOP), 5-methylsulfinylpentyl GS (5MSOP), 7-methylsulfinylheptyl GS (7MSOH), 4-methylthiobutyl GS (4MTB), 8-methylsulfinyloctyl GS (8MSOO), Indol-3-ylmethyl GS (I3M), 4-methoxy-indol-3-ylmethyl GS (4MOI3M), and 1-methoxy-indol-3-ylmethyl GS (1MOI3M). “Total glucosinolates” in Fig. 3 refers to the sum of these compounds.

Morphological responses

The effects of phyB inactivation and MeJA treatment on plant morphology were characterized in 4-week old rosettes using two typical shade avoidance markers: leaf hyponasty (leaf angle with respect to the horizontal plane) and lamina:petiole (L:P) ratio (*10*). Plants were transferred to the corresponding light treatments (Amb or FR) 2 h before the end of the photoperiod on day -1. On day 0, the 7$^{th}$ youngest leaf was identified in each rosette, and all rosettes were sprayed with the indicated concentration of MeJA, 3 h after the beginning of the photoperiod. On day 1, spraying with MeJA was repeated 3 h after the beginning of the photoperiod, and plants were harvested at the end of the photoperiod for morphological measurements. In addition to the standard FR supplementation treatment of 28 µmol $s^{-1}$ $m^{-2}$ (indicated as FRH in this figure), an attenuated FR treatment of 14 µmol $s^{-1}$ $m^{-2}$ was used in these experiments (denoted as FRL).

*Botrytis cinerea* culture and infection bioassays

A suspension of *Botrytis cinerea* (strain B05) spores was used to inoculate individual leaves of 3-week old rosettes, as described previously (*29*). After 48 h, infected leaves (3 per plant) were collected and photographed. Lesion areas were calculated using Adobe Photoshop software (Adobe Systems, USA). In each experiment, there were 14 plants for each "genotype x light treatment" combination (n = 14), and the reported lesion area is the average of the damaged areas of the 3 leaves infected in each plant.

*Spodoptera littoralis* bioassays

Eggs of *Spodoptera littoralis* where supplied by Syngenta Crop Protection (Switzerland). After 2-3 days at 18°C, the eggs hatched and the larvae were fed on an artificial diet based on white bean as described in (*43*). For insect growth bioassays, 5-7-day old caterpillars were placed on 4-5-week old plants under the specified light treatments (3 caterpillars per plant: n = 10 plants per treatment and genotype). After 5 days, caterpillars were collected and weighted. In some experiments, plants were harvested, immediately frozen in liquid nitrogen and stored at -80°C until used for extraction of glucosinolates.

Identification and phylogenetic analysis of ST2a-like proteins

In order to find homologs of ST2a in other species of green plants, the KEGG (http://www.genome.ad.jp/kegg/) nucleotide and protein sequence databases for monocot and dicot genomes were scanned. With these amino acid sequences, a set of ST2a-like proteins represented in 8 monocot and 22 dicot species was completed. Additionally, we included all the *A. thaliana* sulfotransferase (SOT) proteins sequences identified by Piotrowski et al. (*44*) to conduct a global phylogenetic analysis. Identification codes for sequences are listed in Data File S2. The alignment of the full-length amino acid sequences was performed in ClustalW using standard settings (Gonnet weight matrix, gap opening = 10 and gap extension = 0.2) and was adjusted by visual inspection. Bayesian phylogenetic analyses on aligned full-length sequences were performed with MrBayes v. 3.1.2 setting an MCMC algorithm (*45*). The evolutionary distances were computed using the Dayhoff matrix based method and are in the units of the number of amino acid substitutions per site. The rate variation among sites was modeled with a gamma distribution. The analysis involved 48 amino acid sequences. Three independent runs were computed for 1 000 000 generations. All trees were visualized using the program MEGA 5.01 (*46*).

Statistical analyses

Data from real time PCR, metabolites, morphological measurements and bioassays were analyzed using factorial analysis of variance (ANOVA). When the interaction term was significant ($P < 0.05$), differences between means were tested using a post-hoc Tukey test. Appropriate transformations of the primary data were used when needed to meet the assumptions of the ANOVA. Statistical analyses were carried out using INFOSTAT software (*47*).

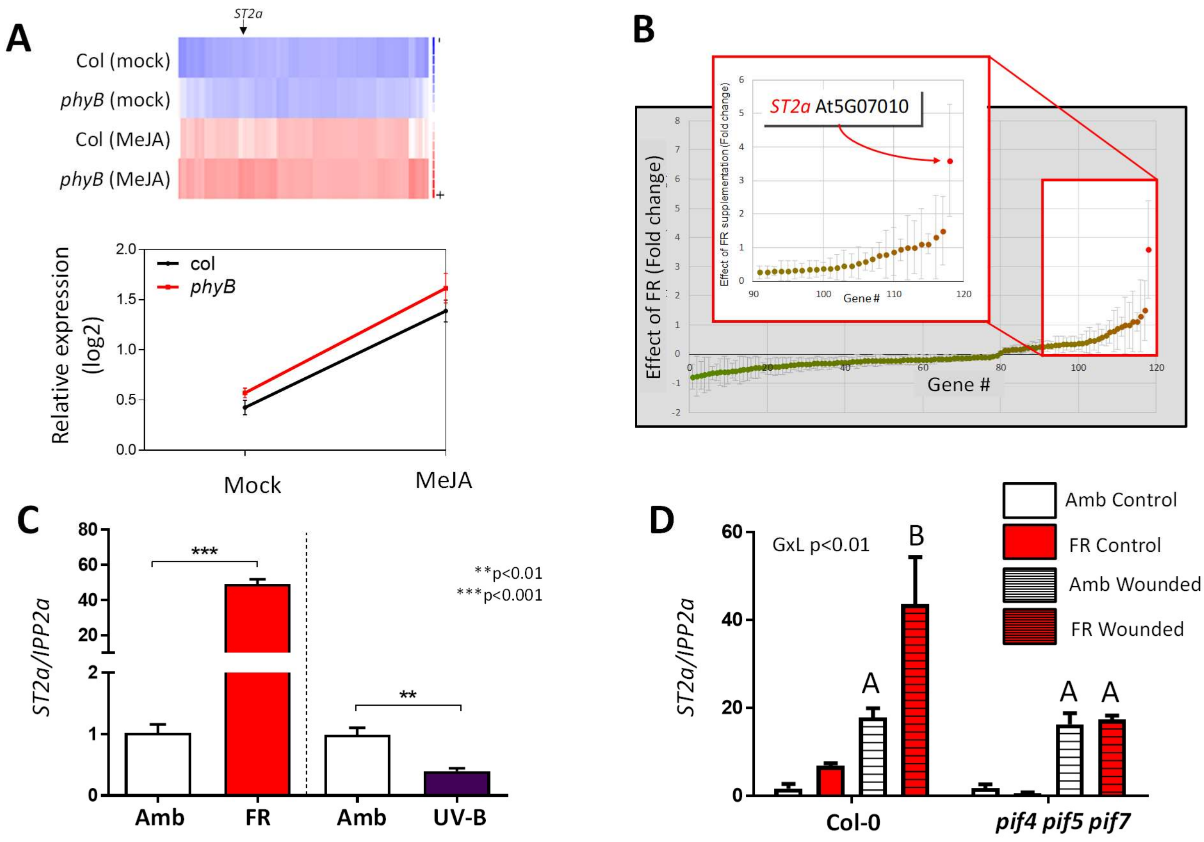

**Fig. S1. AT507010 (*ST2a*) belongs to a small cluster of MeJA-upregulated genes that are also upregulated under conditions of phyB inactivation, via the phyB-PIF module.** (A) Heat map (upper part) showing a cluster of genes (approximately 25 % of the genes upregulated by 200 µM MeJA) that are expressed more strongly in the *phyB-9* mutant than in Col-0 plants, and mean expression values (lower part) of this set of genes. This cluster contains AT5G07010 (*ST2a*). (B) Examination of publicly available microarray data shows that among the genes included in categories with GO terms that contain the word "jasmonic", AT5G07010 stands out for being the most strongly up-regulated gene in plants exposed to low R:FR ratios (see arrow). *ST2a* was also reported to be upregulated by continuous FR radiation (*48*). For each gene, dots indicate the average of 7 independent microarray experiments [see details in ref. (*37*)]; bars indicate 95 % confidence intervals. (C) qPCR analysis demonstrated strong upregulation of *ST2a* in response to supplemental FR radiation and downregulation by supplemental UV-B radiation under our experimental conditions (5-week-old Col-0 plants; relative expression data were normalized to the mean of the Ambient control; samples were taken 5 h (FR) or 6 h (UV-B) after the start of the radiation supplementation treatment). (D) In a *pif4pif5pif7* triple mutant, which does not activate morphological responses to low R:FR ratios, the transcription of *ST2a* was normally upregulated by mechanical wounding, but not at all by supplemental FR radiation (3-week-old plants). **, p<0.01; ***, p<0.001.

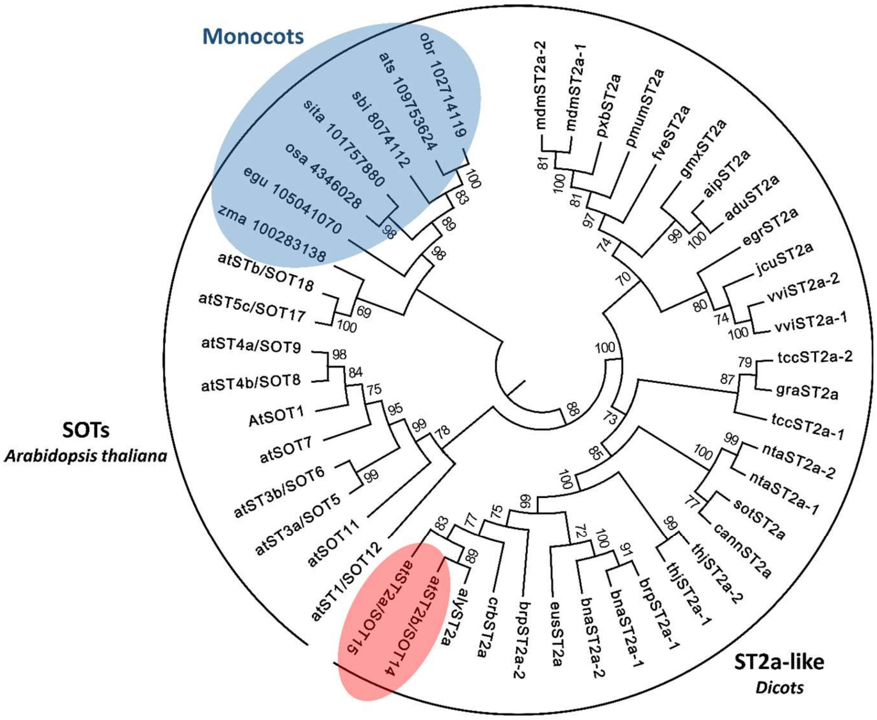

*adu  Arachis duranensis*
*aip  Arachis ipaensis*
*aly  Arabidopsis lyrata*
*ats  Aegilops tauschii*
*bna  Brassica napus*
*brp  Brassica rapa*
*cann Capsicum annuum*
*crb  Capsella rubella*
*egr  Eucalyptus grandis*
*egu  Elaeis guineensis*
*eus  Eutrema salsugineum*
*fve  Fragaria vesca*
*gmx  Glycine max*
*gra  Gossypium raimondii*
*jcu  Jatropha curcas*
*mdm  Malus domestica*
*nta  Nicotiana tabacum*
*obr  Oryza brachyantha*
*osa Oryza sativa japonica*
*pmum Prunus mume*
*pxb  Pyrus x bretschneideri*
*sbi  Sorghum bicolor*
*sita Setaria itálica*
*sot  Solanum tuberosum*
*tcc Theobroma cacao*
*thj  Tarenaya hassleriana*
*vvi  Vitis vinifera*
*zma  Zea mays*

**Fig. S2. Phylogeny of Arabidopsis sulfotransferases (SOT).** Bayesian phylogenetic analyses on aligned full-length sequences were performed with MrBayes v. 3.1.2 setting an MCMC algorithm (*45*). The evolutionary distances were computed using the Dayhoff matrix based method, and are in the units of the number of amino acid substitutions per site (see Materials and Methods for details).

| Enzyme | Function | FR effect on gene |
|---|---|---|
| ILL6 / IAR3 | Amido-hydroxylation of JA-Ile and 12OH-JA-Ile | ↑ |
| JOX2 & JOX4 | Hydroxylation of JA | ↑(*JOX2*) ↓(*JOX4*) |
| CYP94B3 | Hydroxylation of JA-Ile | ↑ |
| JAR1 | Forms JA-amido conjugates, including JA-Ile | Not affected by MeJA or FR (p>0.05) |
| ST2a | Sulfation of OH-JA *in vitro* | ↑↑↑ |
| CYP94C1 | Carboxylation of JA-Ile | Not affected by FR (p>0.05) |

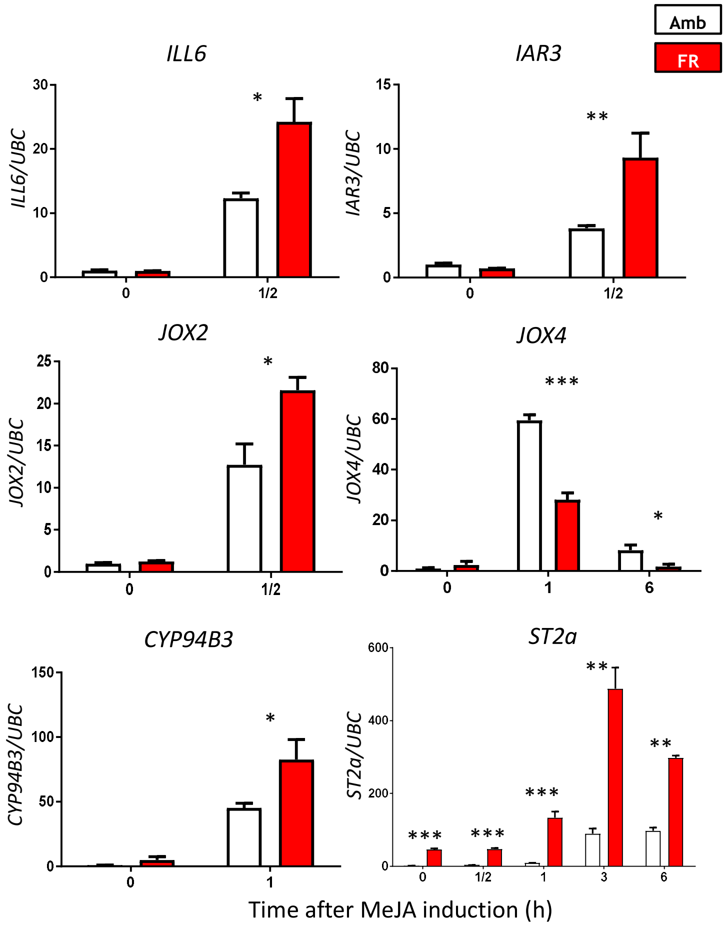

*p<0.05
**p<0.01
***p<0.001

**Fig. S3. FR regulates the transcription of several genes involved in jasmonate metabolism.** Expression of genes encoding for the following enzymes were measured by qPCR in Col-0 Arabidopsis rosettes exposed to two light conditions (Amb or FR) at different times (0, 0.5, 1, 3 and 6 h) after MeJA (200 uM) treatment: ILL6/IAR3 (*49*); JOX/JAO (*27, 50*); CYP94B3 and CYP94C1 (*51-53*); ST2a (*21*); and JAR1 (*54, 55*). The bar charts show the relative expression data, only for those genes and time points in which FR had a significant (p<0.05) effect. Thin bars indicate 1 SE.

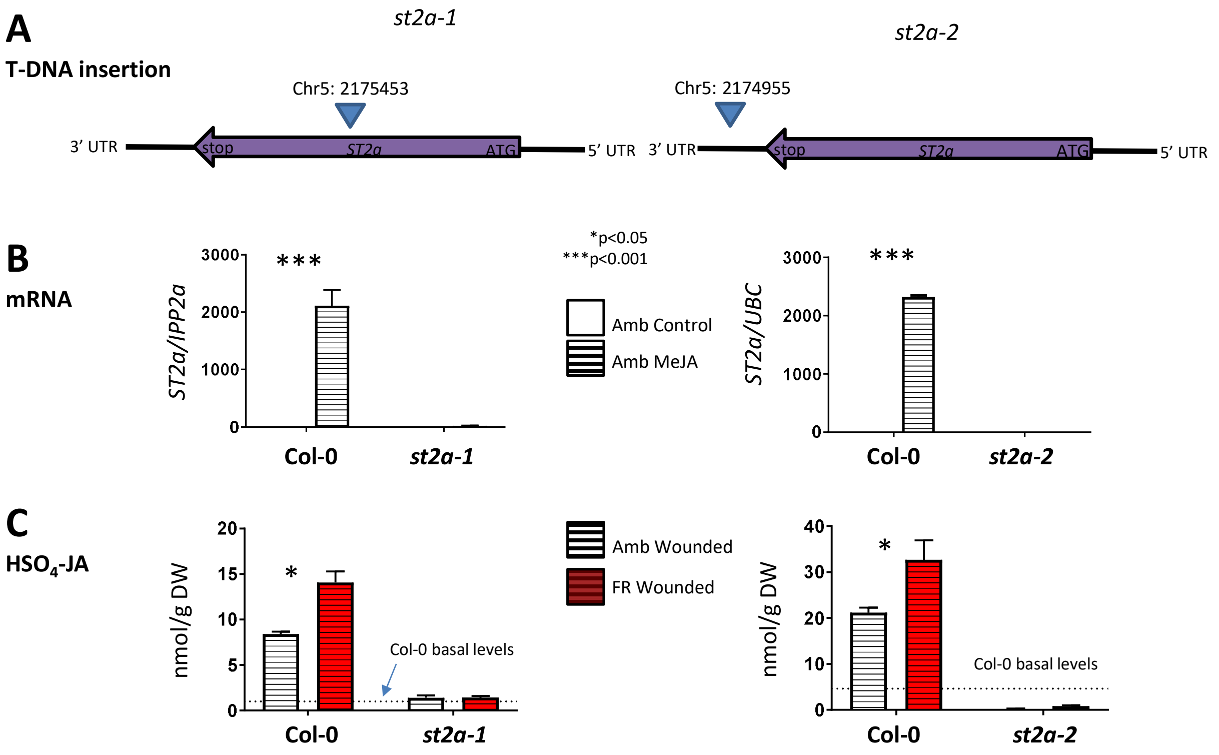

**Fig. S4. *st2a-1* and *st2a-2* are *ST2a* null mutants, and *ST2a* is required for the production of $HSO_4$-JA in vivo.** T-DNA position, *ST2a* mRNA levels, and $HSO_4$-JA concentrations in two different *st2a* null mutants*:* *st2a-1* (GABI_149G04) and *st2a-2* (SALK_ 075656). (A) Representative scheme of *ST2a* gene, showing coordinates for T-DNA insertion. (B) *ST2a* gene expression showed strong regulation by MeJA (200 µM, 4 h after treatment) in wild type (Col-0) plants, and it was nearly undetectable in both *st2a* mutant lines. (C) Both *st2a* mutant lines failed to accumulate $HSO_4$-JA after wounding (4 h) under ambient or FR light conditions.

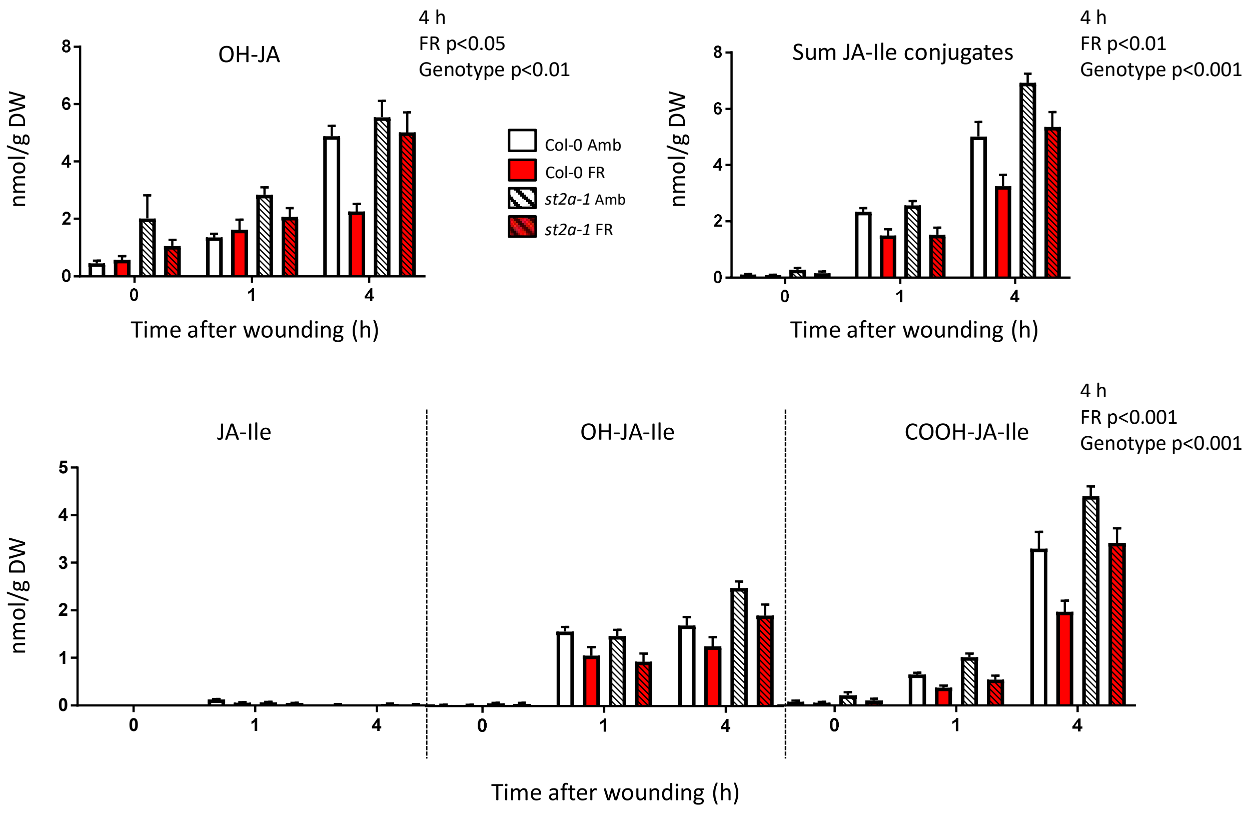

**Fig. S5. FR radiation reduced the concentrations of OH-JA and the sum of JA-Ile conjugates in wounded plants and the concentration of these compounds was higher in *st2a-1* than in Col-0.** Significant ($P < 0.05$) terms in the factorial analysis are indicated for each panel. Thin bars indicate 1 SE ($n = 6$).

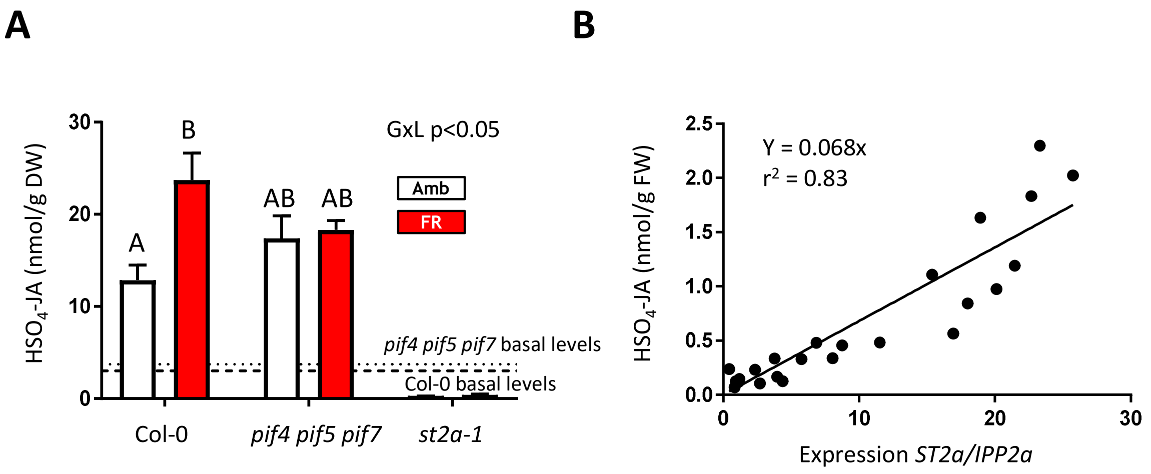

**Fig. S6. The accumulation of $HSO_4$-JA is determined by the expression of *ST2a*.** A) FR fails to upregulate the pool of $HSO_4$-JA in a *pif4 pif5 pif7* triple knock-out mutant, where FR fails to upregulate *ST2a* expression. Samples for metabolite analysis were taken 4 h after wounding (n = 6 independent pools of 3 rosettes each; thin bars indicate 1 SE). The dotted/dashed lines parallel to the abscissa indicate the basal levels of $HSO_4$-JA in plant of both genotypes harvested before the wounding treatment. B) In Col-0 plants, there is a tight, significant correlation between *ST2a* mRNA and $HSO_4$-JA concentration in rosette tissue. Different levels of *ST2a* gene expression were obtained by varying the amount of mechanical damage (see Materials and Methods for details) and intensity of FR supplementation (between 14 and 45 µmol $s^{-1}$ $m^{-2}$ of lateral FR). Samples were taken 4 h after wounding (each datum point represents an independent pool of 3 rosettes).

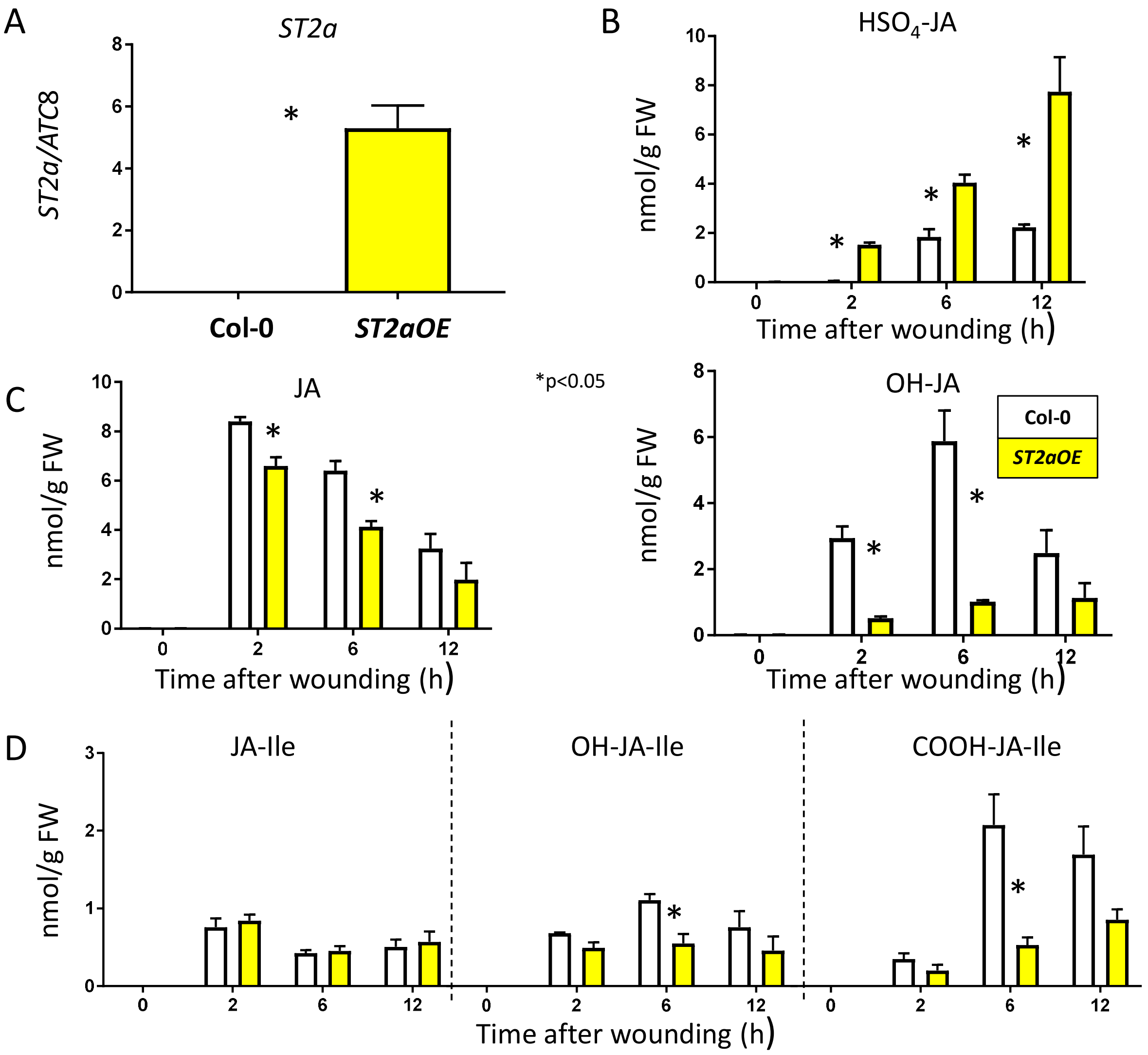

**Fig. S7. Overexpression of *ST2a* increases $HSO_4$-JA at the expense of other JA metabolites.** (A) Constitutive *ST2a* mRNA accumulation in the leaves of *ST2aOE* compared to Col-0. (B-D) Time course of $HSO_4$-JA and other indicated JA metabolite accumulation in wounded leaves of Col-0 and *ST2aOE*. Asterisks denote significant difference ($p<0.05$, Student's *t* test) compared to Col-0 at each given time points. Thin bars indicate 1 SE (n = 3 biological replicates). gFW, gram fresh weight.

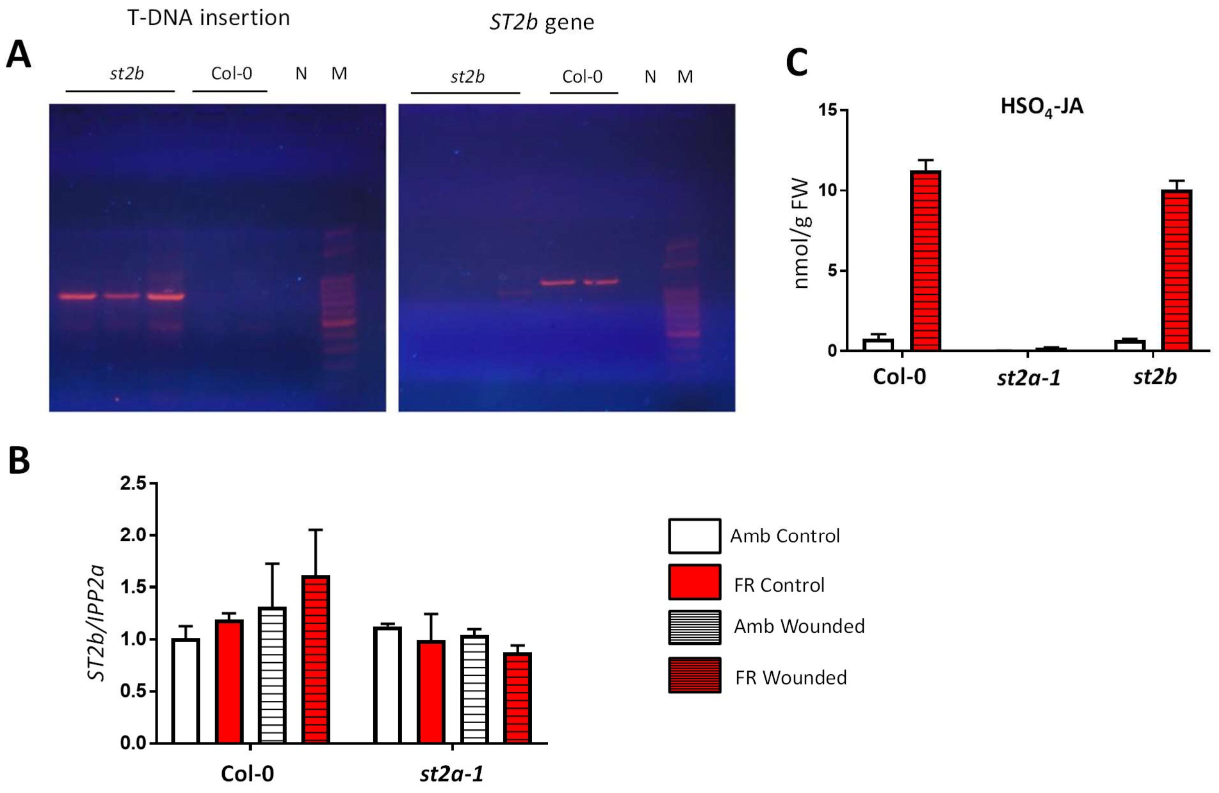

**Fig. S8. A *st2b* knock out mutant has normal concentrations of $HSO_4$-JA. (**A) Genotyping of a *st2b* null mutant. Images show photographs of PCR products run on an agarose gel (TAE buffer, agarose 1.3%). The primers were designed to amplify the T-DNA insertion (Left photograph, showing PCR product only in *st2b* mutants) or DNA flanking the region of the *ST2b* gene where the T-DNA is inserted (Right photograph, showing PCR product only in Col-0, as the interruption of this region by the T-DNA (10 kb) is too long to produce product in *st2b* mutant). N = negative control. M = ladder. (B) *ST2b* gene expression is not induced by wounding or FR radiation, and it is not altered by the *st2a-1* mutation (n = 3 pools of three individual rosettes). Rosettes of 3-week old plants were exposed to the indicated light treatments (Amb or FR); wounding was performed by pressing with a forceps every mature leaf. Plants were harvested before (control) or after wounding (4 h) for gene expression analysis. (C) A *st2b* null mutant produces wild type levels of $HSO_4$-JA. Plants of the indicated genotypes were exposed to the FR treatment and harvested 4 h after wounding for phytohormone analysis (n = 6 pools of three individual rosettes). Primers used of qPCR or genotyping are listed in Table S1.

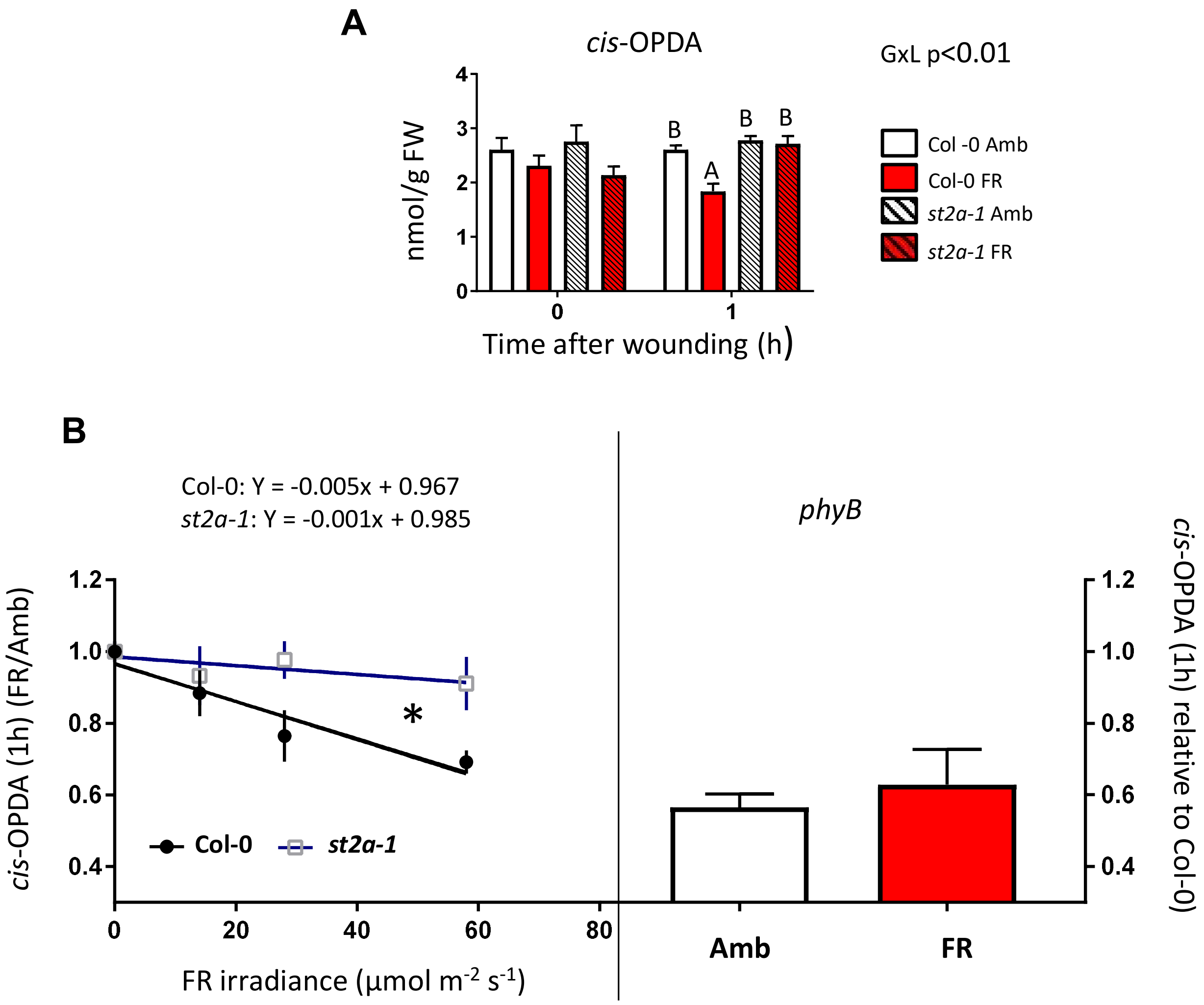

**Fig. S9. Phytochrome B inactivation reduces the accumulation of *cis*-OPDA in Col-0, which is consistent with the down-regulation of *LOX2* gene expression in Col-0.** (A) cis-OPDA concentration in Arabidopsis rosettes 1 h after wounding. The significance of the genotype x light (GxL) interaction term is indicated. Different letters indicate significant ($P < 0.05$) differences between means; thin bars indicate 1 SE ($n = 6$ biological replicates). (B) Relationship between the relative concentration of OPDA (FR/Amb) 1 h after wounding in Col-0 and *st2a-1* rosettes and the irradiance of FR received by the plants. Thin bars indicate 1 SE; the asterisk denotes a significant ($P < 0.05$) difference between the slopes of the fitted regression lines. The right panel indicates the relative OPDA concentrations in plants of the *phyB-9* mutant, which was used as a control for phyB inactivation (FR irradiance 28 µmol $s^{-1}$ $m^{-2}$).

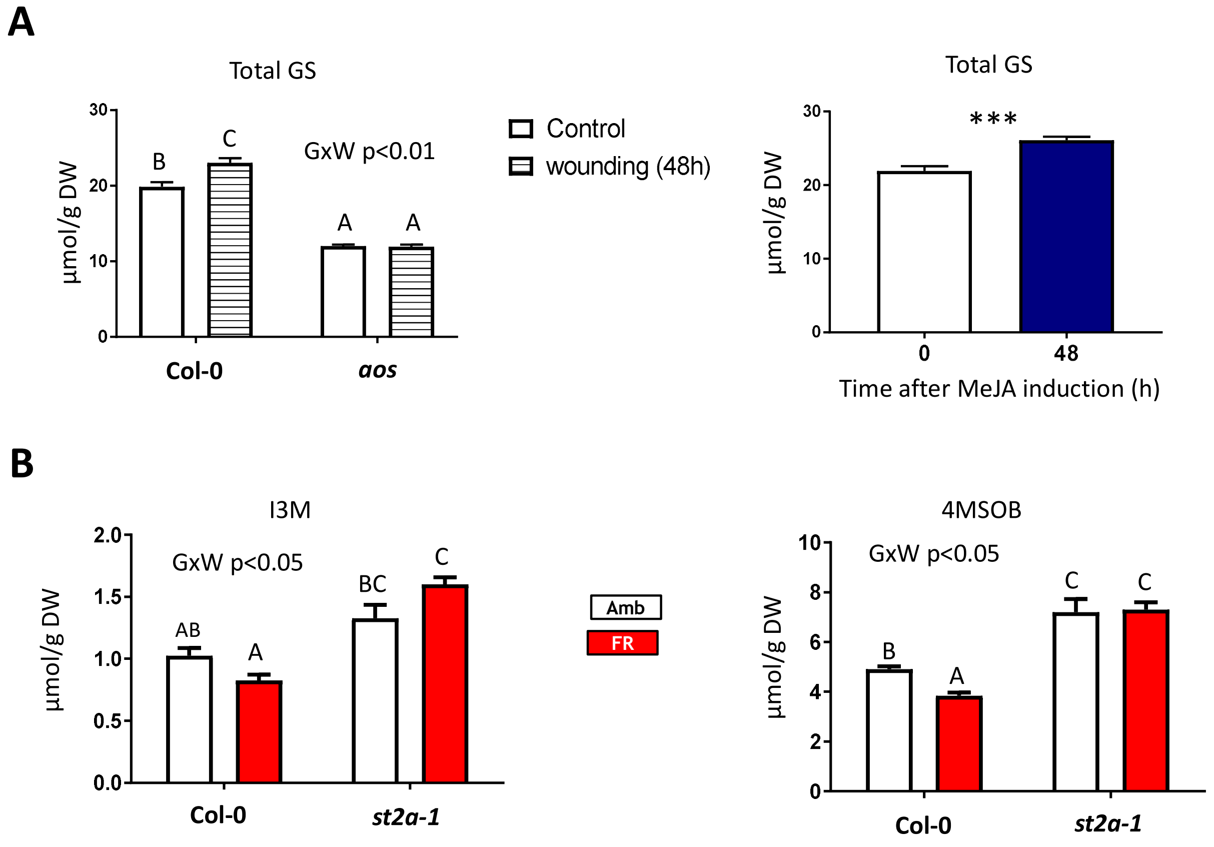

**Fig. S10. Glucosinolate concentrations in induced plants**. (A) Glucosinolate accumulation requires JA synthesis to respond to wounding (left panel), and it is inducible by treatment with exogenous MeJA (right panel). (B) Effects of FR on accumulation of indolic (I3M, indol-3-ylmethyl) and aliphatic (4MSOB4-methylsulfinylbutyl) glucosinolates in wounded Col-0 and *st2a-1* plants. Samples were taken 48 h after wounding or MeJA treatment (200 μM). When comparing light responses between genotypes, the significance of the genotype x light (GxL) interaction term is indicated. Different letters indicate significant ($P < 0.05$) differences between means; thin bars indicate 1 SE (n = 6 biological replicates).

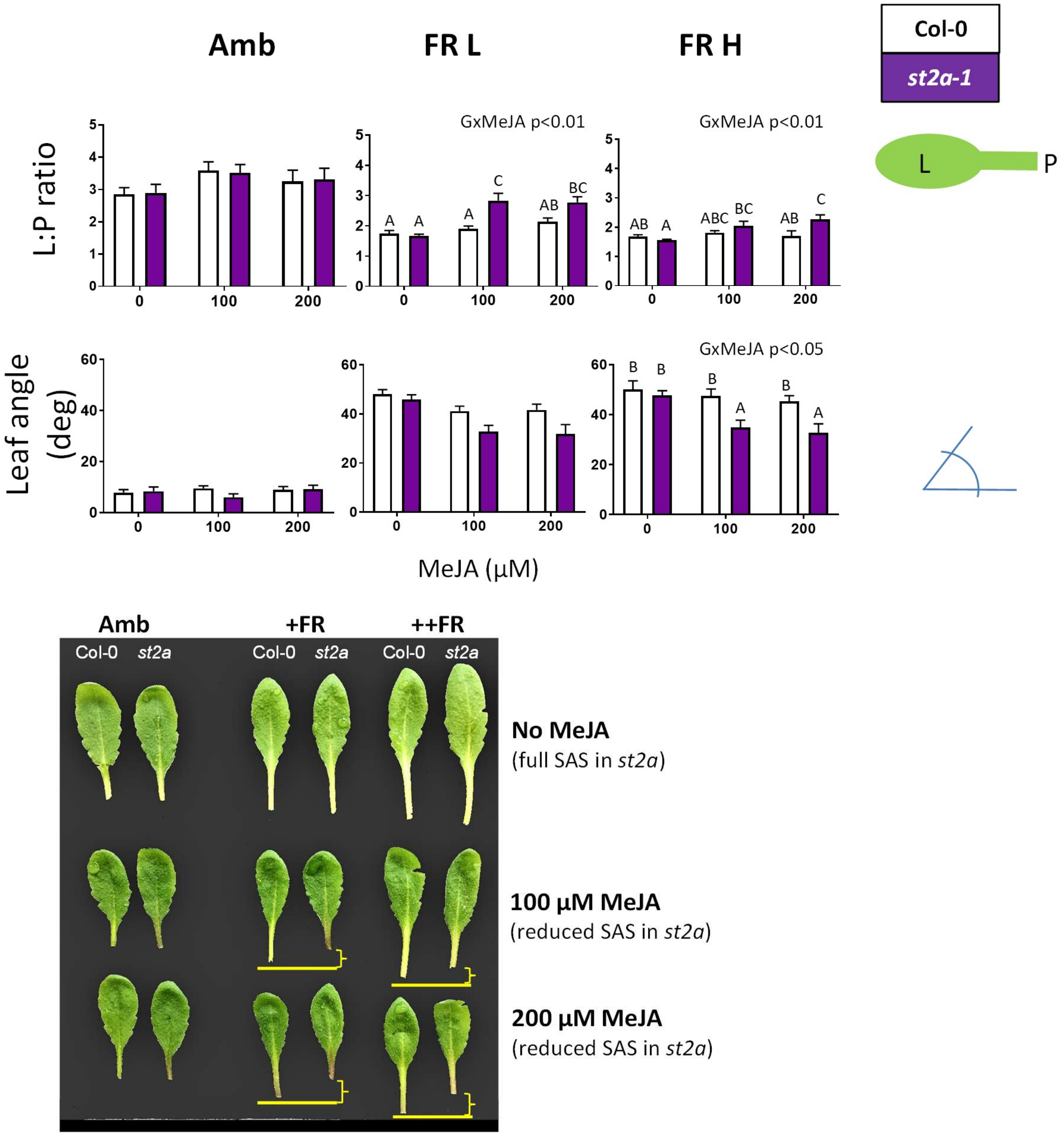

**Fig. S11. *ST2a* is required for the full expression of shade avoidance responses to low R:FR ratio**. Col-0 rosettes responded to supplemental FR radiation with a reduction in the lamina:petiole (L:P) ratio (upper panel) and increased leaf hyponasty (middle panel). *st2a-1* rosettes had normal phenotypes under control conditions but, compared with Col-0 rosettes, they displayed impaired shade-avoidance responses when exposed to low doses of MeJA. Pictures of representative leaves are shown in the lower panel. FRL = 14 µmol $s^{-1}$ $m^{-2}$; FRH = 28 µmol $s^{-1}$ $m^{-2}$.

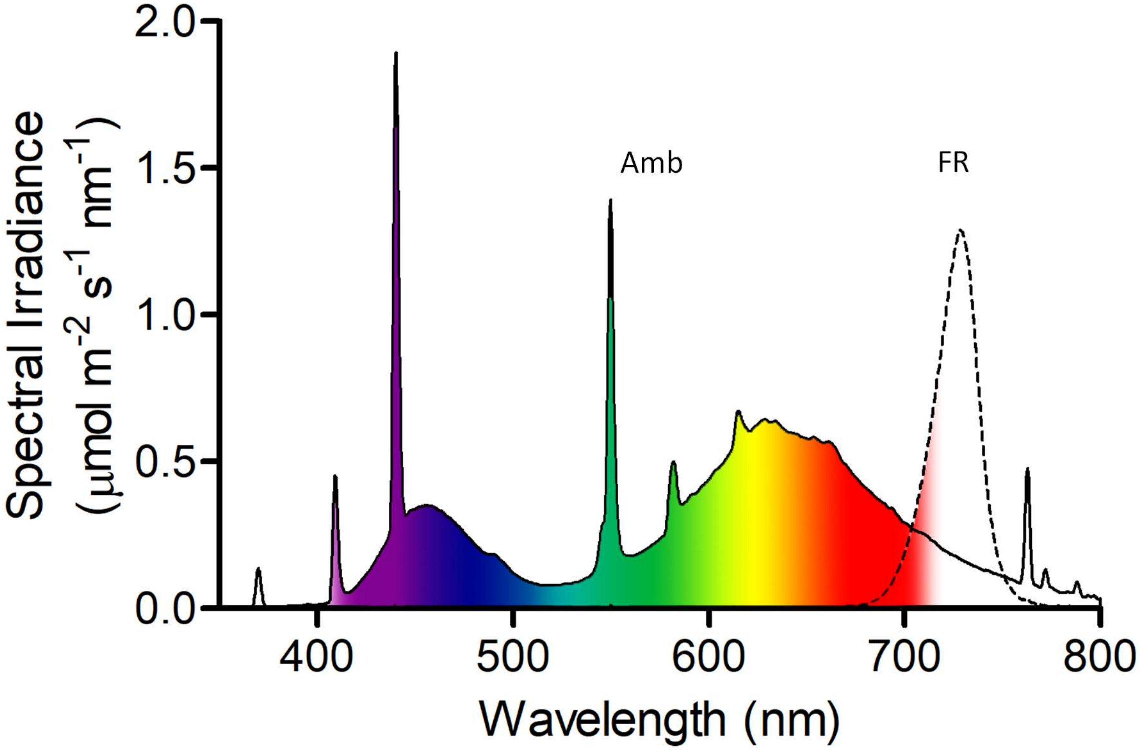

**Fig. S12. Spectral scans of the light sources used in the experiments.** Amb = "Ambient" light (110 µmol $s^{-1}$ $m^{-2}$), provided by fluorescent bulbs (solid line); FR = supplement of lateral FR radiation (dotted line) added during the course of the photoperiod to plants of the FR treatment (except indicated otherwise, the FR irradiance was 28 µmol $s^{-1}$ $m^{-2}$. Photoperiod was always 10 h.

**Table S1. Sequences of primers used in genotyping or qPCR**

| **Genotyping** | |
|---|---|
| Primer name | Sequence 5' to 3' |
| st2a-1 Fw | AAAGTTCTTGATCGACTTGT |
| st2a-1 Rv | AACATTTCCAATCCCTCG |
| st2a-2 Fw | CACAAGTGGAAAGATTGTCAG |
| st2a-2 Rv | TAATCATTGTGGTTCAGTCTC |
| st2b Fw | ATTGCTCAACAACCCCCTC |
| st2b-Rv | ATTCGGTCGAGAATCCCAG |
| **qPCR** | |
| *IPP2a* Fw | ATGGTTCAGATTGGTGGTGGAC |
| *IPP2a* RV | AAAGATGTTCAGAGTTTGTGGATGG |
| *UBC* Fw | CTGCGACTCAGGGAATCTTCTA |
| *UBC* Rv | TTGTGCCATTGAATTGAACCC |
| *ST2a* Fw | CTGAGGGCCTACTATATACG |
| *ST2a* Rv | CGACAAACTTCGGTGTTGAC |
| *MYC2* Fw | CCGAAAACCCGAATCTGGAT |
| *MYC2* Rv | GGGTCTGAGAATGAACCGGAC |
| *LOX2* Fw | AAGACTGACCAGCGGATTACG |
| *LOX2* Rv | CAGGCATCTCAAACTCGCAC |
| *VSP2* Fw | TGACCGTTGGAAGTTGTGGA |
| *VSP2* Rv | CGAACCATTAGGCTTCAATATGAG |
| *IAR3* Fw | GATGCACTTGCTATGCAGGA |
| *IAR3* Rv | ACACTCCAGCCTCCACAATC |
| *ILL6* Fw | AGGCATTGTATCCCGTGAAG |
| *ILL6* Rv | CGGGTATATCGCATTCTGCT |
| *JOX2* Fw | CGGCGAAGAGCTAGTGAAGC |
| *JOX2* Rv | TGGTCATACCGCCAGGATCG |
| *JOX4* Fw | GAGGAGGCGACAAAGTCGGA |
| *JOX4* Rv | CATCACACACGATGGACCTGA |
| *CYP94B3* Fw | TGGCTTACACGAAGGCTTGTC |
| *CYP94B3* Rv | AGTCCCACGAAACTGGAGGAT |
| *JAR1* Fw | TCACGCTTTTAGAACCTTTGAACAG |
| *JAR1* Rv | GGACCGATGGGACAGTAATACG |
| *CYP94C1* Fw | GGCCCGGATTACGAAGAGTTT |
| *CYP94C1* Rv | GGCAACTTACCTTCGTT |
| *ST2b* Fw | GATCCAGAACTATGAGAACCGG |
| *ST2b* Rv | CTGAAAGTGAGACCAGATCCAG |

**Table S2. Details of analysis of phytohormones by LC-MS/MS [HPLC 1260 (Agilent Technologies)-QTRAP6500 (SCIEX)] in negative ionization mode**

| Q1 | Q3 | RT (min) | Compound | Internal std | RF | DP | EP | CE | CXP |
|---|---|---|---|---|---|---|---|---|---|
| 136.93 | 93 | 3.3 | SA | D4-SA | 1.0 | -20 | -8 | -24 | -7 |
| 263 | 153.2 | 3.4 | ABA | D6-ABA | 1.0 | -20 | -12 | -22 | -2 |
| 209.07 | 59 | 3.6 | JA | D6-JA | 1.0 | -20 | -9 | -24 | -2 |
| 322.19 | 130.1 | 3.9 | JA-Ile | D6-JA-Ile | 1.0 | -50 | -4.5 | -30 | -4 |
| 290.9 | 165.1 | 4.6 | OPDA | D6-JA | 1.0 | -20 | -12 | -24 | -2 |
| 263 | 165 | 4.2 | dinor-OPDA | D6-JA | 0.7 | -20 | -10 | -20 | -10 |
| 338.1 | 130.1 | 3 | OH-JA-Ile | D6-JA-Ile | 1.0 | -50 | -4.5 | -30 | -4 |
| 225.1 | 59 | 2.6 | OH-JA | D6-JA | 1.0 | -20 | -9 | -24 | -2 |
| 352.1 | 130.1 | 3 | COOH-JA-Ile | D6-JA-Ile | 1.0 | -50 | -4.5 | -30 | -4 |
| 305 | 97 | 2.4 | SulfoJA | D6-JA | 6.0 | -20 | -10 | -60 | -10 |
| 387.1 | 207 | 2.4 | JA-Gluc | D6-JA | 3.7 | -50 | -10 | -28 | -21 |
| 140.93 | 97 | 3.3 | D4-SA | | | -20 | -8 | -24 | -7 |
| 269 | 159.2 | 3.4 | D6-ABA | | | -20 | -12 | -22 | -2 |
| 215 | 59 | 3.6 | D6-JA | | | -20 | -9 | -24 | -2 |
| 214 | 59 | 3.6 | D5-JA | | | -20 | -9 | -24 | -2 |
| 328.19 | 130.1 | 3.9 | D6-JA-Ile | | | -50 | -4.5 | -30 | -4 |
| 327.19 | 130.1 | 3.9 | D5-JA-Ile | | | -50 | -4.5 | -30 | -4 |

++++++++++++

**Data File S1. Overrepresented GO categories in RNAseq data reported on Fig 3B (Excel File).**
**Data File S2. Identification codes for sulfotransferase sequences (excel File).**

++++++++++++
